# Leaf Damage Is Associated with Alteration of Microbiome Composition and Abundance in the *Spartina alterniflora* Phyllosphere

**DOI:** 10.64898/2026.04.21.719896

**Authors:** Jose L. Rolando, Anna L. Carnes, Michael Hodges, Heather Joesting, Joel E. Kostka

**Affiliations:** Microbiology and Cell Science Department, Institute of Food and Agricultural Sciences (IFAS). Fort Lauderdale Research and Education Center, University of Florida. Davie, Florida, USA; School of Biological Sciences, Georgia Institute of Technology, Atlanta, Georgia, USA; South Carolina Department of Natural Resources, Charleston, South Carolina, USA; Department of Biology, Georgia Southern University, Savannah, Georgia, USA; School of Earth and Atmospheric Sciences, Georgia Institute of Technology, Atlanta, Georgia, USA; Center for Microbial Dynamics and Infection, Georgia Institute of Technology, Atlanta, Georgia, USA

## Abstract

*Spartina alterniflora*, the dominant plant species in salt marshes along the Atlantic and Gulf of Mexico coastlines of the Americas, is affected by disease and sudden vegetation dieback. Despite the foundational role of *S. alterniflora* in low-elevation salt marshes, the response of the native leaf-associated microbiome (i.e., phyllosphere microbiome) to leaf damage resulting from disease and environmental stress has not been explored. We hypothesized that healthy and damaged plants would show differentiation in their phyllosphere microbiomes following primary infection or exposure to environmental stressors. Here, we analyzed changes in prokaryotic and fungal relative abundance, diversity, and community composition in the *S. alterniflora* phyllosphere microbiome. We compared natural marsh and greenhouse plants in Georgia and South Carolina, USA, and collected leaves from healthy and damaged natural plants across two contrasting *Spartina* phenotypes that differ in their exposure to environmental stress. Our results show that plant origin (i.e., greenhouse vs. natural marsh), plant health status (i.e., healthy vs. damaged), and plant phenotype (i.e., short vs. tall *Spartina*) affect microbial relative abundance, alpha diversity, and community composition in the *S. alterniflora* phyllosphere. Damaged leaves presented higher microbial abundance and alpha diversity than healthy leaves, suggesting microbial proliferation following leaf damage. Plants raised from seeds in the greenhouse presented the lowest microbial abundance and Shannon diversity for both prokaryotic and fungal communities, indicating that in natural ecosystems the phyllosphere microbiota is acquired predominantly through horizontal transmission from the environment. Overall, this study provides novel insights into the assembly of the *S. alterniflora* phyllosphere microbiome.

**Importance:** Salt marshes are tidally influenced coastal wetlands that provide a range of ecosystem services to global and local communities, including protection from storm surge, water purification, and carbon sequestration. *Spartina alterniflora* is the dominant plant species in Atlantic and Gulf of Mexico marshes in the Americas. Fungal disease and exposure to environmental stressors have previously been described in marsh ecosystems and linked to extensive and sudden vegetation dieback in the southeastern U.S. In this study, we propose that microbial proliferation follows plant damage caused by either fungal disease or environmental stress, leading to profound changes in native leaf-associated microbiota abundance, diversity, and composition. Using greenhouse plants as a control, we also demonstrate that microbes colonizing marsh leaves are acquired predominantly from the environment. Overall, this study advances the ecological understanding of the *S. alterniflora* leaf microbiome assembly, establishing a baseline to inform future work harnessing plant-associated microbes for ecosystem restoration.

## Introduction

Salt marsh ecosystems are intertidal wetlands found in low-energy wave zones, often located on the landward side of barrier islands. These ecosystems provide a wide range of services, including water purification, recreational fishing, tourism, carbon sequestration, and cultural benefits that support overall human well-being (1). Coastal ecosystems worldwide are under threat and degrading due to environmental stressors such as limited sediment supply, accelerating sea-level rise, and land-use changes (2, 3).

*Spartina alterniflora* is the dominant plant species in low-lying salt marshes native to the Gulf of Mexico and the eastern coast of the Americas (4). Within its distribution range, the health of the marsh is therefore directly linked to the health of *S. alterniflora*. During the past few decades, large-scale vegetation dieback has occurred in the United States, with fungal pathogens identified as one of multiple stressors driving these events (2, 5, 6). Several species of the genus *Fusarium* have been associated with dieback in salt marshes and have been shown to be pathogenic to *S. alterniflora* (6, 7). Besides *Fusarium* spp., fungal species from the *Claviceps* and *Puccinia* genera have also been implicated in disease of *S. alterniflora* (8, 9). However, little is known about how the natural microbial assemblage of the *S. alterniflora* phyllosphere responds to environmental stressors and disease. Studies in model organisms and agricultural plants have shown that microbial infection alters the phyllosphere microbiome, increasing the abundance of opportunistic commensals with an overall reduction in alpha diversity (10–12). This profound change in the diversity and composition of the native phyllosphere microbiome is referred to as dysbiosis and is thought to further deteriorate host health (13).

*Spartina alterniflora* dominates the plant community of low-lying salt marshes due to its high tolerance to salinity and flooding stress (14). One of *S. alterniflora*’s key adaptations is the excretion of salt through specialized leaf glands, which results in the accumulation of salt crystals on the leaf surface (15). The high salt content on leaves creates a unique and selective habitat for microbial colonization. Furthermore, environmental stress gradients exist within *S. alterniflora* stands as a function of distance from large tidal creeks (Figure 1). Plants located farther from tidal creeks tend to accumulate higher concentrations of salts and phytotoxic sulfides in their root zone (16). This environmental gradient results in differentiated phenotypic expression in *S. alterniflora*: stressed plants grow in a stunted short phenotype (<100 cm), whereas less stressed creekhead plants can reach heights of up to 200 cm (17). Limited studies have begun to characterize the microbiome and virome associated with the roots and surrounding sediments of *S. alterniflora* across this salt marsh stress gradient (18–20). However, few studies have examined the microbial communities inhabiting the leaf environment using cultivation-independent techniques (21–23), and their response to leaf damage and environmental stress remains largely unknown.

**Figure 1.**
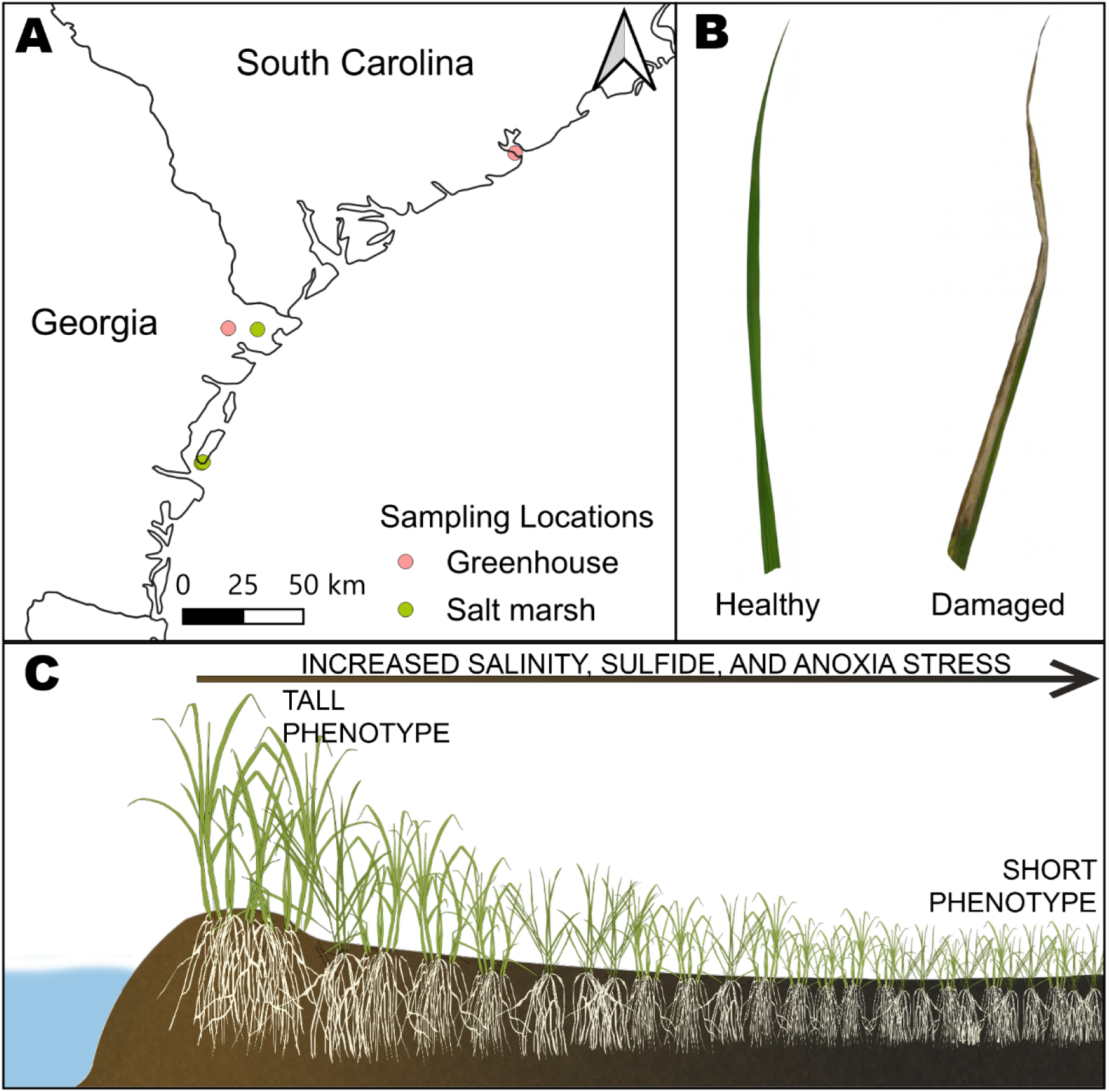
Study site and environmental context for Spartina alterniflora leaf collection. Map of sampling locations along the southeastern U.S. Atlantic coast showing natural salt- marsh sites and greenhouse sampling locations (A). Representative examples of healthy and damaged S. alterniflora leaves used for phyllosphere microbiome analyses (B). Conceptual figure of the natural marsh environmental gradient, illustrating the transition from tall to short S. alterniflora phenotypes associated with increasing salinity, sulfide concentration, and sediment anoxia stress (C). Conceptual figure adapted from Mendelssohn and Morris (2002).

By analyzing changes in prokaryotic and fungal community composition across various plant health states, natural stress gradients, and greenhouse and natural marsh settings, this study aimed to identify the forces driving microbial community assembly in the *S. alterniflora* phyllosphere. The objectives of this study were to: (i) define the *S. alterniflora* phyllosphere microbiome and quantify the effects of plant health and environmental stress on microbial relative abundance, diversity, and composition; (ii) elucidate the source of the *S. alterniflora* phyllosphere microbiome in the southeastern U.S.; and (iii) identify prokaryotic and fungal taxa associated with environmental stress and leaf damage in the *S. alterniflora* phyllosphere. We predicted that plant health, environmental stress, and growth conditions would serve as selective forces in the assembly of *S. alterniflora* phyllosphere microbiomes. This study was designed to test the hypothesis that damaged *S. alterniflora* plants show a dysbiotic state in which a few opportunistic prokaryotic and fungal species dominate the phyllosphere. Additionally, greenhouse plants grown from field-collected botanical seeds (i.e., not vegetative propagules) were used to test the source of the *S. alterniflora* phyllosphere microbiome by differentiating horizontal from vertical transmission during microbiome recruitment.

## Results

### Overview of sampling design

The phyllosphere microbiome was characterized based on leaf health status across two barrier islands in the state of Georgia and two greenhouse settings, one in Savannah, Georgia, and the other in Charleston, South Carolina (Figure 1A). Leaf health status was classified as healthy or damaged based on field observations. Damaged leaves presented one or more of the following: lesions, necrosis, spotting, or visible fungal growth, whereas healthy leaves lacked visible signs of tissue damage (Figure 1B). Only leaves from the top third of the plant were collected to avoid senescing leaves. For natural marsh plants, leaves were collected across a natural stress gradient in sulfide concentration, salinity, and sediment redox conditions (Figure 1C). This environmental gradient results in differentiated phenotypic expression in *S. alterniflora*: stressed plants express a short phenotype (<100 cm), whereas plants in less stressful environments are classified as the tall phenotype (>100 cm) (17). Previous studies have shown that plants of the short phenotype present lower rates of primary production and nutrient uptake and increased root fermentation rates compared to the tall phenotype (16, 17).

A total of 247 leaves were sampled over two years during peak growth of *S. alterniflora* in July 2021 and 2022 (detailed sampling effort in Table 1). Amplicon sequencing of the 16S rRNA gene and ITS region was performed to characterize the composition and diversity of the prokaryotic and fungal phyllosphere communities, respectively (further details in Materials and Methods). Amplicon sequence analysis yielded 3,507,700 and 737,654 total 16S rRNA gene and ITS reads, respectively. A total of 6,629 unique prokaryotic 16S rRNA gene amplicon sequence variants (ASVs), 159 fungal ITS ASVs, and 65 plant ITS ASVs were obtained.

**Table 1.**
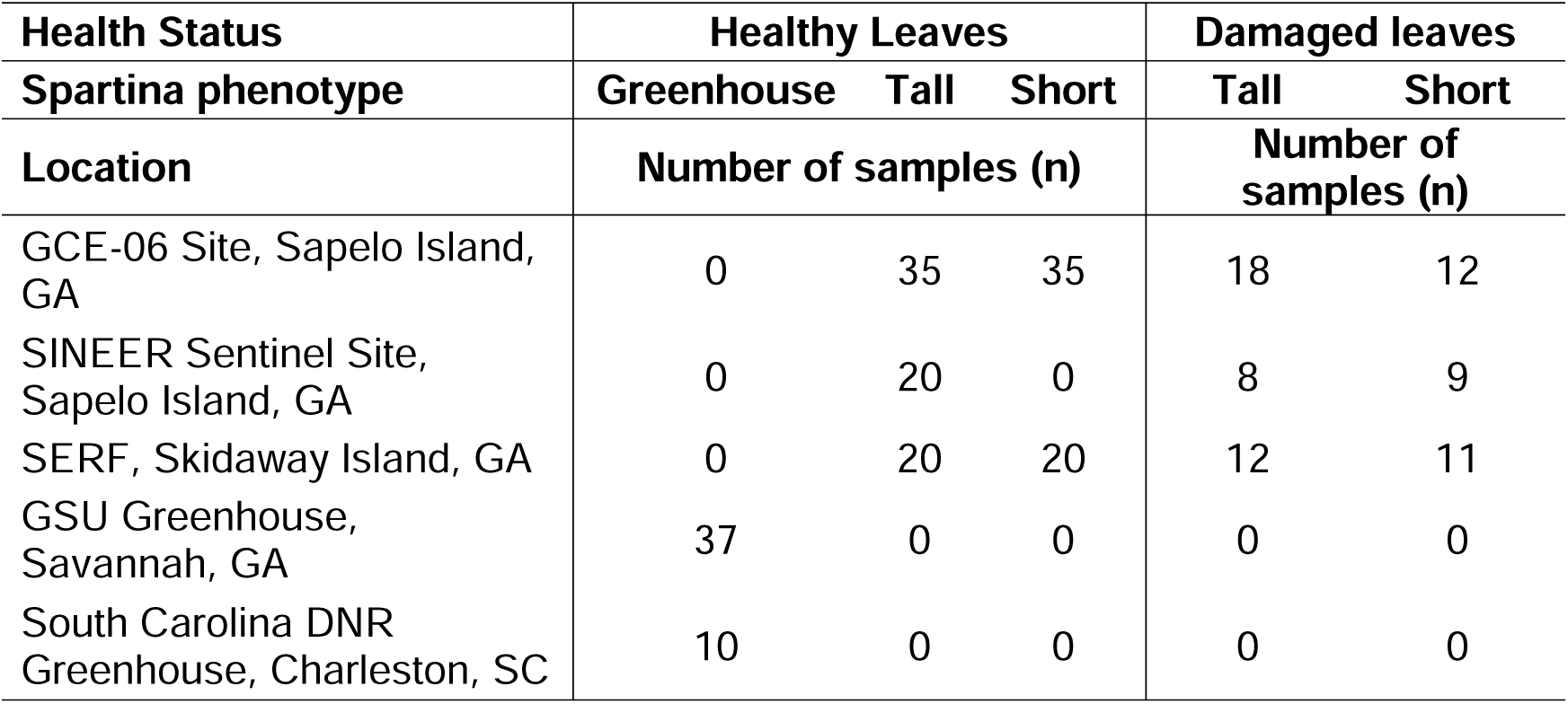
Sampling effort of S. alterniflora leaves by sampling location, leaf health status and plant phenotype.

| Health Status | Healthy Leaves |  |  | Damaged leaves |  |
| --- | --- | --- | --- | --- | --- |
| Spartina phenotype | Greenhouse | Tall | Short | Tall | Short |
| Location | Number of samples (n) |  |  | Number of samples (n) |  |
| GCE-06 Site, Sapelo Island, GA | 0 | 35 | 35 | 18 | 12 |
| SINEER Sentinel Site, Sapelo Island, GA | 0 | 20 | 0 | 8 | 9 |
| SERF, Skidaway Island, GA | 0 | 20 | 20 | 12 | 11 |
| GSU Greenhouse, Savannah, GA | 37 | 0 | 0 | 0 | 0 |
| South Carolina DNR Greenhouse, Charleston, SC | 10 | 0 | 0 | 0 | 0 |

### Leaf damage and environmental stress affect phyllosphere microbiome abundance and diversity

We analyzed changes in microbial relative abundance, diversity, and composition in leaf samples collected from natural salt marshes to assess the effect of leaf damage and environmental stress in situ. We quantified prokaryotic relative abundance as the percentage of prokaryotic 16S rRNA gene ASV reads relative to total 16S rRNA gene sequences, including those from plant organelles (i.e., mitochondria and chloroplasts). In both plant phenotypes, prokaryotic relative abundance was significantly higher in damaged leaves than in healthy leaves (Figure 2A). Similarly, fungal relative abundance, quantified as the percentage of fungal ITS reads relative to total ITS sequences (plant + fungal), was significantly higher in damaged leaves in both phenotypes (Figure 2B). When assessing microbial alpha diversity, damaged leaves from the tall phenotype presented higher prokaryotic Shannon diversity than healthy tall leaves, while no significant differences were observed in the short phenotype (Figure 2C). Fungal Shannon diversity was higher in damaged tall leaves only when compared to healthy short leaves (Figure 2D). However, in both prokaryotic and fungal communities, ANOVA indicated that leaf health status significantly affected Shannon diversity, with an observed increase in alpha diversity in damaged leaves (Table S1).

**Figure 2.**
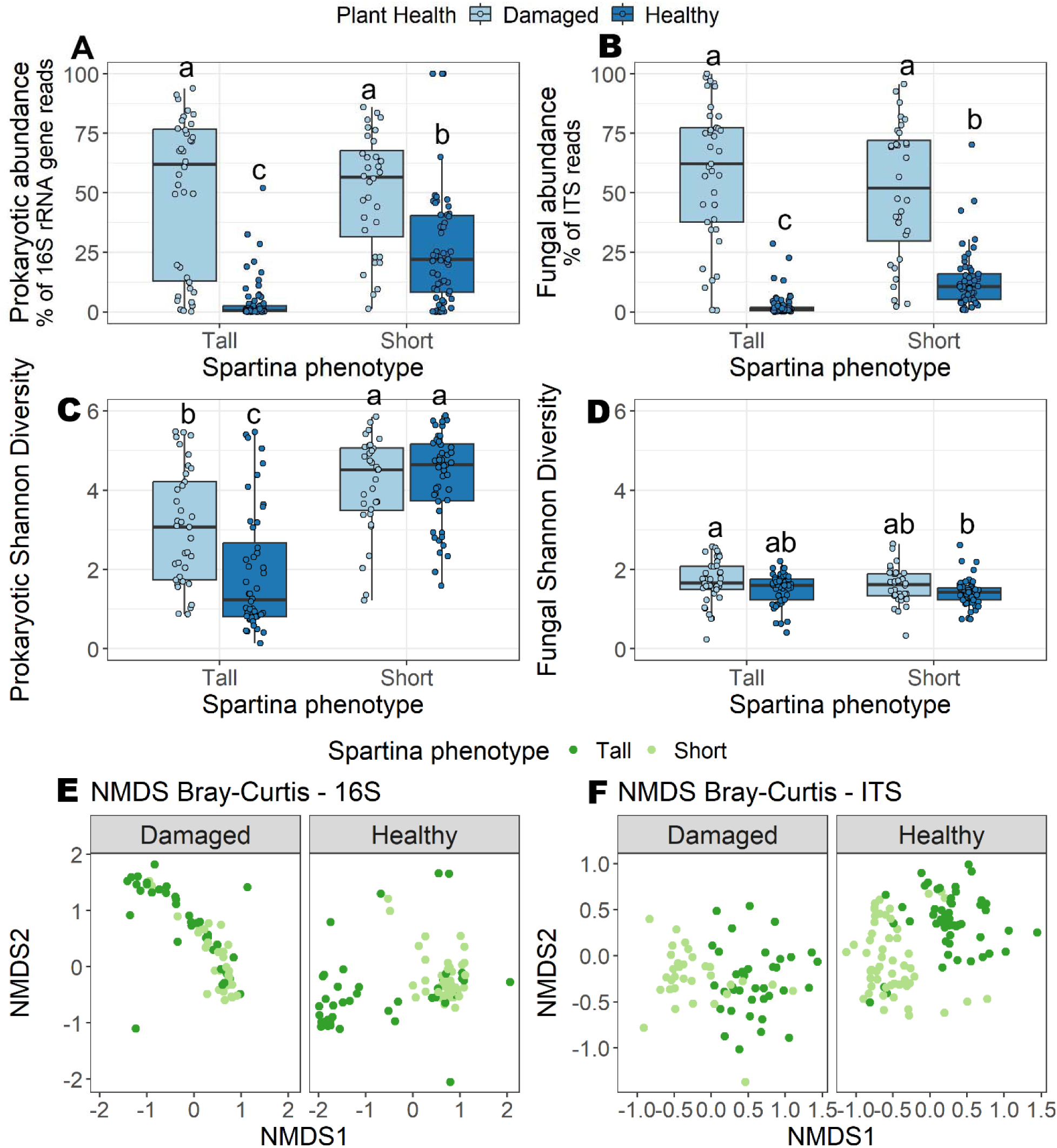
Microbiome abundance and diversity are affected by plant health and environmental stress. Boxplots of prokaryotic (A) and fungal (B) relative abundances by plant health and S. alterniflora phenotype. Boxplots of prokaryotic (C) and fungal (D) alpha diversity measured by the Shannon index per plant health and phenotype. Non-metric multidimensional scaling (nMDS) ordination of the Bray-Curtis dissimilarity matrix of phyllosphere prokaryotic (E) and fungal (F) communities with colors representing plant phenotype. In boxplots, boxes represent the upper and lower interquartile; the median is represented as a horizontal line within the boxes; and whiskers extend to values within 1.5 × IQR. Different letters indicate significant differences (p < 0.05) based on Tukey’s (Abundance; A, B) and Dunn’s (Shannon diversity; C, D) post hoc tests.

Microbial abundance and diversity were also affected by the natural stress gradient. Within healthy leaves, the stressed short phenotype had greater prokaryotic and fungal relative abundance compared to the tall phenotype (Figures 2A, 2B). Further, we observed higher prokaryotic phyllosphere alpha diversity in the short phenotype in both damaged and healthy leaves compared to the tall phenotype (Figure 2C). No plant phenotype effect was observed on fungal alpha diversity (Figure 2D).

To identify the primary factors shaping prokaryotic and fungal phyllosphere community composition in natural *S. alterniflora* plants, we performed PERMANOVA on Bray-Curtis dissimilarity matrices (Table 2). As evidenced by both PERMANOVA and NMDS ordination analysis, prokaryotic community composition was primarily affected by plant health status, followed by plant phenotype and site location (Figure 2E, Table 2). In turn, fungal community composition was primarily affected by plant phenotype, followed by health status, and least by site location (Figure 2F, Table 2).

**Table 2.**
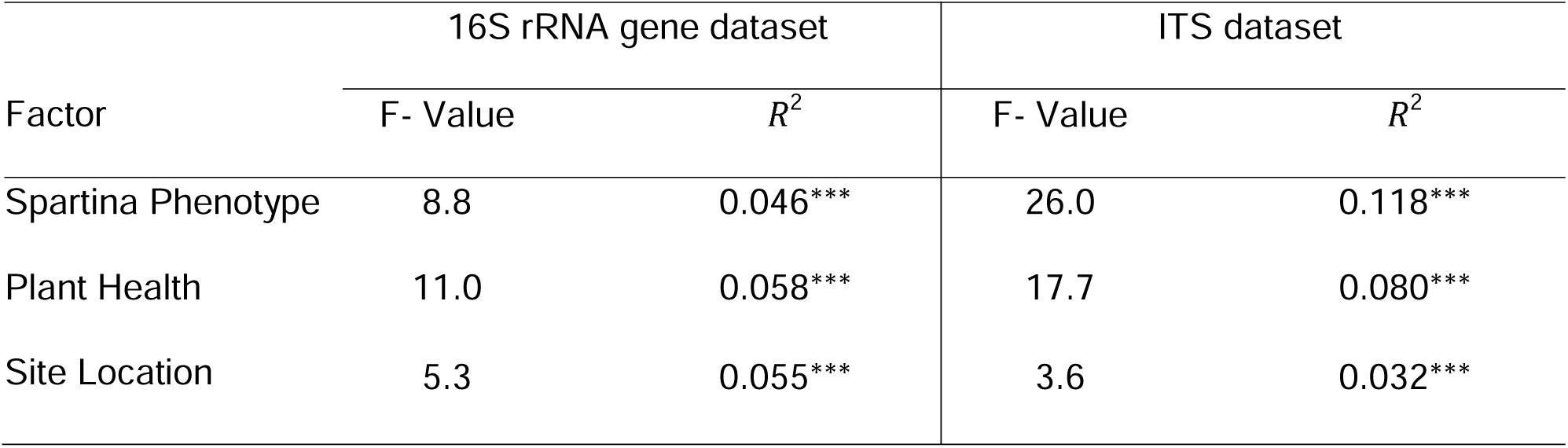
PERMANOVA analysis of factors affecting the community composition of the phyllosphere microbiome of S. alterniflora for prokaryotes and fungi in natural ecosystems using the Bray-Curtis metric with 999 permutations. *** represents p-value < 0.001.

| Factor | 16S rRNA gene dataset |  | ITS dataset |  |
| --- | --- | --- | --- | --- |
| | F- Value | $R^2$ | F- Value | $R^2$ |
| Spartina Phenotype | 8.8 | 0.046*** | 26.0 | 0.118*** |
| Plant Health | 11.0 | 0.058*** | 17.7 | 0.080*** |
| Site Location | 5.3 | 0.055*** | 3.6 | 0.032*** |

### Greenhouse plants showed lower microbial abundance and diversity and a distinct phyllosphere microbiome

Leaf samples were collected from greenhouse-grown plants in Savannah, GA, and Charleston, SC, propagated from botanical seeds (i.e., not from vegetative propagules). Seeds were not sterilized so that they retained their natural microbiota upon germination. Since greenhouse plants did not show leaf damage, we compared their phyllosphere microbiome to that of healthy, naturally growing plants from both the short and tall *S. alterniflora* phenotypes. Because greenhouse cultivation for restoration purposes does not replicate the environmental conditions experienced by plants in situ (e.g., tidal flushing, anoxia, biotic interactions with invertebrates and microbes), we did not classify greenhouse plants as either short or tall phenotype; instead, we considered them a distinct phenotypic expression. Phyllosphere prokaryotic and fungal relative abundance was significantly lower in greenhouse plants compared to the two natural *S. alterniflora* phenotypes (Figures 3A, 3B). Median prokaryotic and fungal relative abundances were near zero in greenhouse leaves, with only 0.04% and 0.32% of sequences classified as prokaryotic and fungal, respectively. Further, greenhouse plants had lower prokaryotic diversity than the short phenotype (Figure 3C). In contrast, the greenhouse phyllosphere microbiome showed lower fungal diversity relative to leaves of both the tall and short phenotypes (Figure 3D). Furthermore, NMDS ordination and PERMANOVA based on Bray-Curtis dissimilarity matrices showed that the greenhouse phyllosphere microbiome was compositionally distinct from that of natural plants in both prokaryotic and fungal communities (Figures 3E, 3F, Table 3).

**Figure 3.**
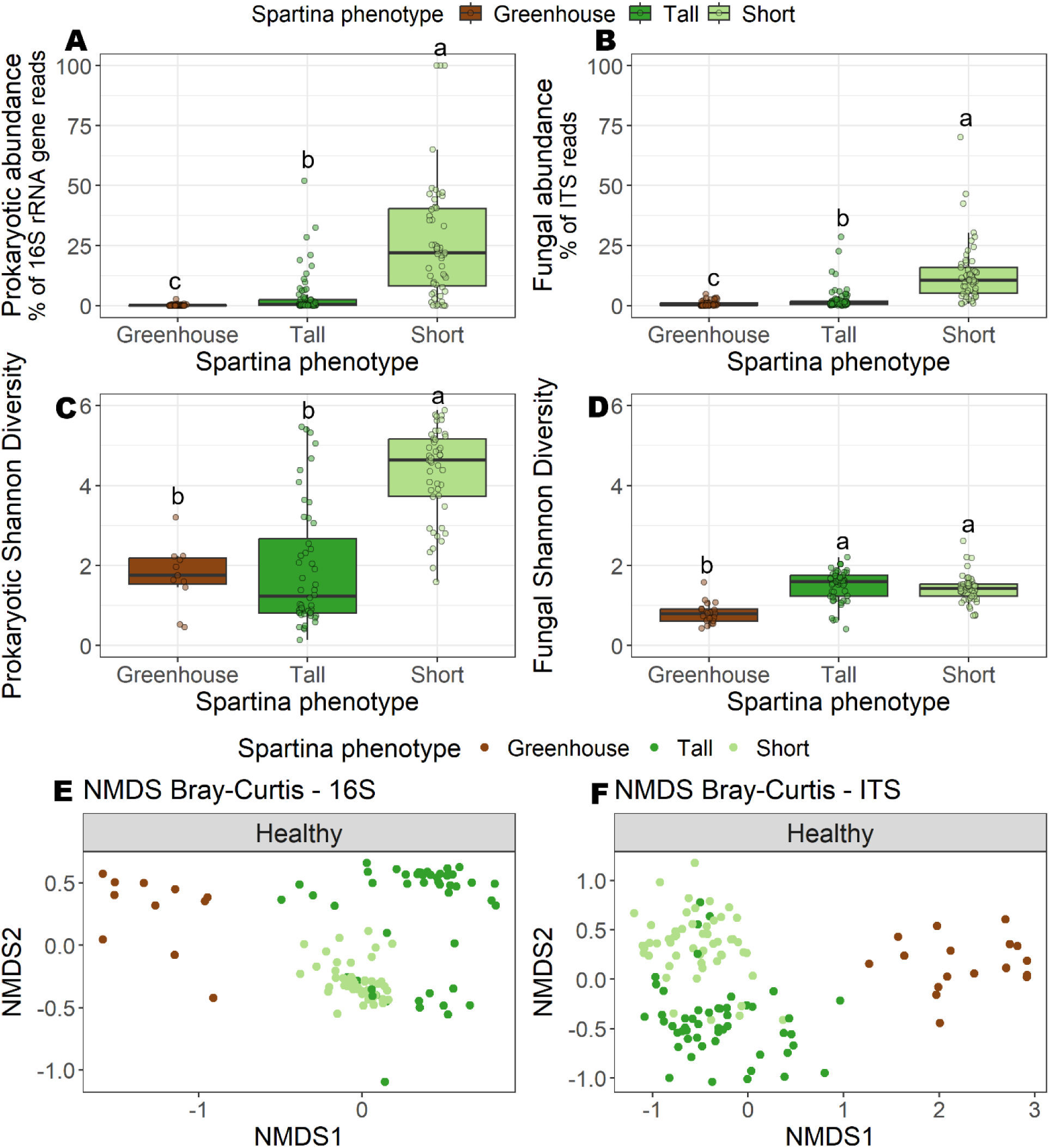
Greenhouse plants have low leaf microbial abundance and diversity. Boxplots of phyllosphere prokaryotic (A) and fungal (B) relative abundances from healthy S. alterniflora plants per plant phenotype and greenhouse-grown plants. Boxplots of phyllosphere prokaryotic (C) and fungal (D) Boxplots of Shannon diversity index from healthy S. alterniflora plants per plant phenotype and greenhouse-grown plants. Non-metric multidimensional scaling (nMDS) ordination of the Bray-Curtis dissimilarity matrix of phyllosphere prokaryotic (E) and fungal (F) communities with colors representing plant phenotype and greenhouse-grown plants. In boxplots, boxes represent the upper and lower interquartile; the median is represented as a horizontal line within the boxes; and whiskers extend to values within 1.5 × IQR. Different letters indicate significant differences (p < 0.05) based on Tukey’s (Abundance; A, B) and Dunn’s (Shannon diversity; C, D) post hoc tests.

**Table 3.** PERMANOVA analysis of factors affecting the community composition of healthy phyllosphere microbiomes of S. alterniflora from short phenotype, tall phenotype and greenhouse plants (Spartina Phenotype) using the Bray-Curtis metric with 999 permutations. *** represents p-value < 0.001.

| Factor | 16S rRNA gene dataset |  | ITS dataset |  |
| --- | --- | --- | --- | --- |
| | F- Value | $R^2$ | F- Value | $R^2$ |
| Spartina Phenotype | 8.4 | 0.066*** | 23.4 | 0.119*** |
| Site Location | 4.3 | 0.067*** | 3.6 | 0.036*** |

### Taxonomic identification of microbes enriched in damaged and stressed leaves

The *S. alterniflora* phyllosphere microbiome in the southeastern USA was composed primarily of bacteria from the Proteobacteria and Bacteroidota phyla and fungi from the Ascomycota phylum (Figures S1, S2). Natural marsh plants presented high relative abundance of bacteria from the *Candidatus* Uzinura, *Kushneria*, *Erythrobacter*, *Salinicola*, and *Salinimonas* genera and fungi from the *Phaeosphaeria*, *Penicillium*, *Haematomma*, and *Cladosporium* genera (Figures S1, S2).

To identify microbial taxa enriched by leaf damage, plant phenotype, and plant origin (i.e., greenhouse vs. natural marsh), we performed differential abundance analysis at the ASV level for both prokaryotic and fungal communities using the Microbiome Multivariable Associations with Linear Models 2 (MaAsLin2; (24)) R package. This analysis was based on total relative abundance, including plant-derived 16S rRNA gene (organelle) and ITS sequences, to assess changes in prokaryotic abundance rather than relative shifts within the microbial community after removal of host sequences. The top 10 microbial ASVs based on lowest q-values were selected for interpretation of microbial differential abundance under different plant growth conditions. Damaged leaves presented increased abundance of selected prokaryotic ASVs from the Gamma- and Alphaproteobacteria classes. ASVs from the genera *Kushneria*, *Pseudoroseicyclus*, *Altererythrobacter*, *Stakelama*, and *Kineococcus* were enriched in damaged *S. alterniflora* leaves (Figure 4A). Greenhouse leaves were in all cases depleted in ASV abundance compared to natural plants (Figure 4B). Within healthy plants, the stressed short phenotype was enriched in ASVs from unclassified Cyanobacteriia and the *Erythrobacter* and *Palleronia*–*Pseudomaribius* genera (Alphaproteobacteria) (Figure 4B), whereas the tall phenotype was enriched only in a Bacteroidia ASV from the *Candidatus* Uzinura genus (Figure 4B). Among damaged plants, the tall phenotype was enriched in ASVs from *Sphingomonas* (Alphaproteobacteria), *Curtobacterium*, *Kineococcus* (Actinobacteria), and an unclassified Gammaproteobacterium, whereas the short phenotype was enriched in ASVs from unclassified Alphaproteobacteria, *Erythrobacter*, *Palleronia*–*Pseudomaribius* (Alphaproteobacteria), and *Salinimonas* (Gammaproteobacteria) (Figure 4C).

**Figure 4:**
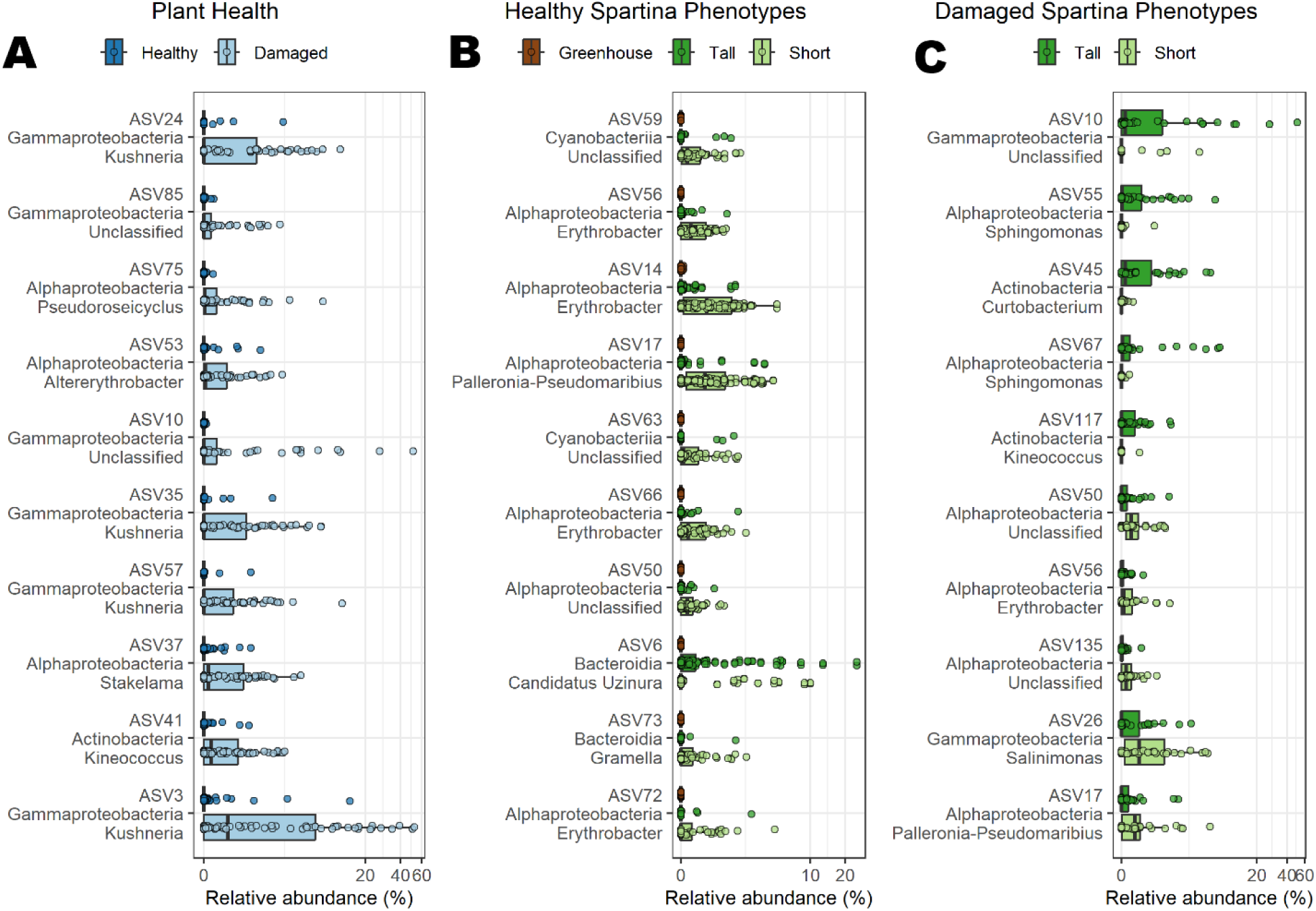
Differential abundance analysis of prokaryotic taxa by plant health and phenotype. Relative abundance boxplots of the 10 most significant fungal amplicon sequence variants (ASVs) associated with plant health (A) and with S. alterniflora phenotype in healthy (B) and damaged (C) plants, as identified by MaAsLin2 (24). Relative abundance values include reads from plant organelle 16S rRNA gene ASVs to reflect a proxy of total ASV abundance. In boxplots, boxes represent the upper and lower interquartile; the median is represented as a horizontal line within the boxes; and whiskers extend to values within 1.5 × IQR.

Differential abundance analysis showed that a fungal ASV from the *Phaeosphaeria* genus (ASV5) was significantly more abundant in damaged leaves, as well as in the phyllosphere of the short phenotype under both healthy and damaged conditions (Figures 5A, B, C). BLAST alignment of ASV5 against the NCBI core nucleotide (nt) database revealed a perfect match of ASV5’s ITS sequence to that of *Phaeosphaeria spartinicola* isolates. Additionally, two further *P. spartinicola* ASVs, a *Cladosporium cladosporioides* ASV, a *Haematomma* ASV, and two ASVs unclassified at the genus level showed increased abundance in damaged leaves compared to healthy plants (Figure 5A). In what appears to be a misannotation in the UNITE taxonomic database, the putative *Haematomma* ASV (ASV8) had a perfect match with several *Papiliotrema mangalensis* strains in the NCBI nt database. Differential abundance analysis of damaged leaves revealed that fungal communities in the tall phenotype were enriched in ASVs from the *Papiliotrema mangalensis*, *Cladosporium cladosporioides*, *Aureobasidium*, *Penicillium*, and *Puccinia* genera (Figure 5C). Damaged leaves from the short phenotype were enriched in ASVs from *Phaeosphaeria spartinicola* and ASVs unclassified at the genus level (Figure 5C).

**Figure 5:**
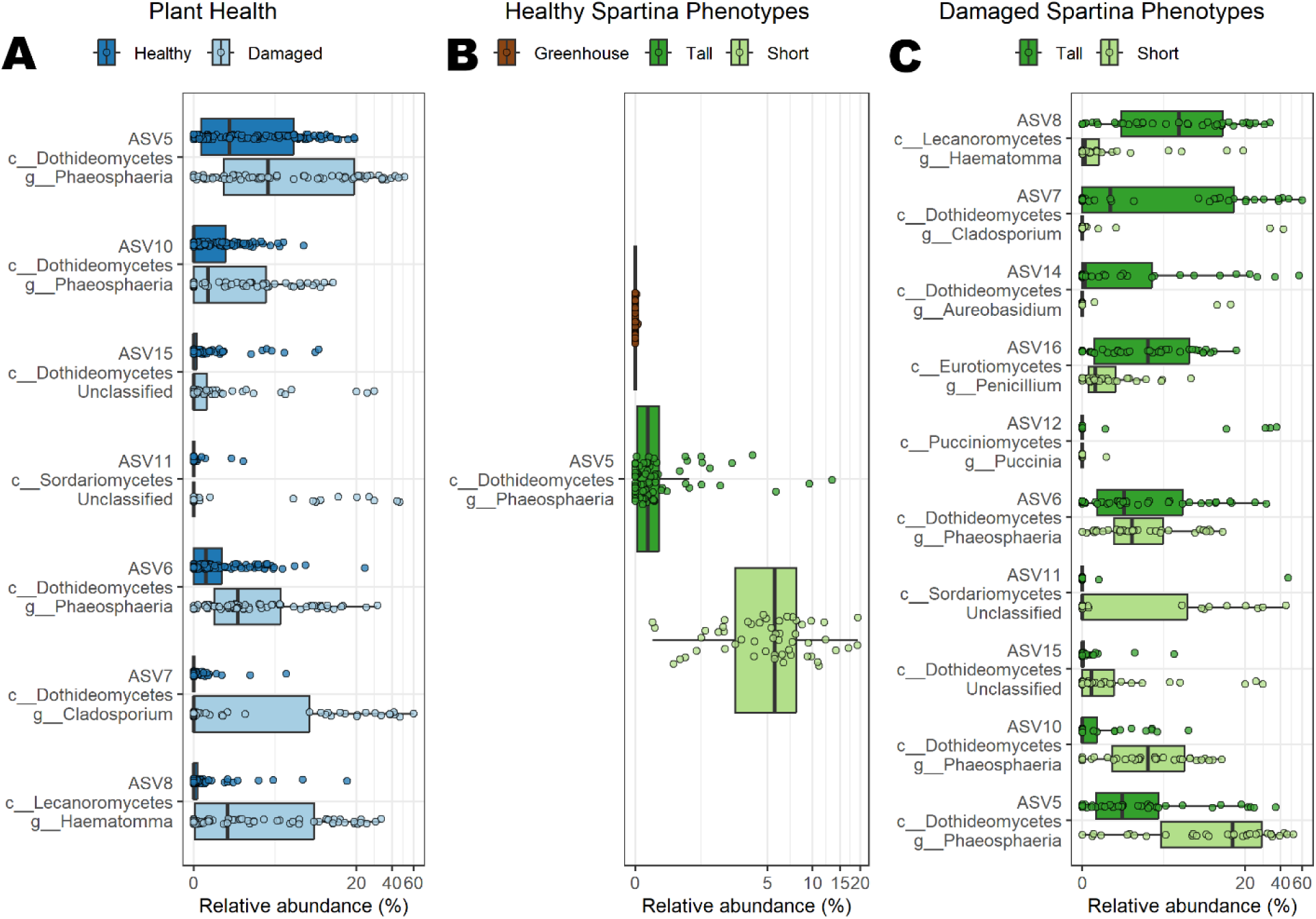
Differential abundance analysis of fungal taxa by plant health and phenotype. Relative abundance boxplots of the most significant prokaryotic amplicon sequence variants (ASVs) associated with plant health (A) and with S. alterniflora phenotype in healthy (B) and damaged (C) plants, as identified by MaAsLin2 (24). Relative abundance values include reads from plant ITS ASVs to reflect a proxy of total ASV abundance. In boxplots, boxes represent the upper and lower interquartile; the median is represented as a horizontal line within the boxes; and whiskers extend to values within 1.5 × IQR.

## Discussion

Here, we defined the *S. alterniflora* phyllosphere microbiome in the southeastern U.S. and, for the first time, examined how environmental stress and leaf health shape its assembly. In contrast to what is commonly found in the phyllosphere of most studied agricultural and model plant species, the most abundant bacterial genera in the *S. alterniflora* phyllosphere were taxa with known adaptations for halophilic growth, including *Salinisphaera*, *Salinicola*, *Salinimonas*, *Kushneria*, and *Erythrobacter* (25–29). This indicates that the phyllosphere microbiome of *S. alterniflora*, in addition to harboring adaptations to common phyllosphere stressors such as oligotrophy, high UV radiation, and low water availability (30), is also adapted to withstand the high-salinity conditions resulting from *S. alterniflora* leaf salt extrusion (15). Salinity has also been shown to shape the phyllosphere microbiome on the salt-secreting leaves of the mangrove *Avicennia marina* (31). In contrast, the most abundant and prevalent fungal species in the *S. alterniflora* phyllosphere was *Phaeosphaeria spartinicola*, a well-known ascomycetous saprobe that mediates lignocellulose degradation of *S. alterniflora* (32). We show that *P. spartinicola* is not only present in decaying plant tissue but is also a prevalent member of the *S. alterniflora* phyllosphere microbiome, including on apparently healthy leaves. Furthermore, we showed that leaf damage and environmental stress trigger profound changes in the phyllosphere of *S. alterniflora* and that the phyllosphere microbiome is acquired predominantly from the surrounding environment, as discussed below.

### Leaf damage and environmental stress are associated with profound changes in the phyllosphere microbiome of S. alterniflora

Microbiome dysbiosis is a concept that originated in the study of the human gut microbiome, in which blooms of pathogenic or harmful bacteria, loss of commensal or beneficial microorganisms, and loss of alpha diversity were observed in association with disease or poor metabolic health (33). Similarly, initial studies of plant microbiomes have found that microbial infections result in an increase in the abundance of opportunistic pathogens while reducing phyllosphere alpha diversity (10–12). Here, we show that leaf damage, whether due to environmental stress or primary fungal infection, causes a profound shift in the *S. alterniflora* phyllosphere microbiome, characterized by an increase in microbial abundance, a shift in microbial composition, and—in contrast to previous studies of phyllosphere dysbiosis—an increase in microbial alpha diversity. The higher alpha diversity observed in damaged leaves from the less stressed tall *Spartina* phenotype is most likely explained by an expansion of phyllosphere niche heterogeneity following leaf damage. Consistent with classical niche and habitat-heterogeneity theory (34, 35), we suggest that mechanical disruption of leaf tissue releases a diversity of carbon substrates and nutrients, increasing resource heterogeneity and facilitating colonization by a wider range of microbial taxa. Increased alpha diversity following leaf damage is more consistent with niche expansion than with pathogen bloom, suggesting that leaf damage in *S. alterniflora* may not follow the same ecological trajectory as dysbiosis described in other systems. Thus, although we observed a profound change in the composition and abundance of microbes following plant damage, we could not confirm further decline in host health due to microbiome dysbiosis.

The higher abundance of *Phaeosphaeria spartinicola* in damaged and stressed leaves must be considered in the context of its interaction with the periwinkle snail (*Littoraria irrorata*). Previous work in *Spartina* marshes has found that lesions from grazers, such as the periwinkle snail, increase leaf decomposition by the saprobic fungus *P. spartinicola* (36, 37). *P. spartinicola* has been found associated with decaying leaves of S. alterniflora in the Atlantic, Gulf of Mexico, and Pacific coasts of the U.S. (21, 38, 39). The periwinkle snail cultivates and harvests *P. spartinicola* growing on *S. alterniflora* leaves as a food source (40). Consistently, we found higher abundance of *P. spartinicola* in damaged leaves, especially within the stressed short phenotype, where periwinkle snail density is the highest (19). This pattern suggests that grazing damage facilitates fungal colonization of the phyllosphere, a process to which *S. alterniflora* is known to be particularly susceptible (41). Thus, high abundance of *P. spartinicola*, or even of periwinkle snails themselves, may serve as an indicator of declining marsh health in *S. alterniflora*-dominated systems. Further studies are needed to better understand the role of the periwinkle snail in shaping the phyllosphere microbiome of *S. alterniflora* and its interactive effects on overall plant physiological status.

*Fusarium* spp. have been described as widespread endophytic and pathogenic fungi *in S. alterniflora*, with implications for sudden-vegetation dieback (7). In particular, *Fusarium palustre* was shown to be present in most *S. alterniflora* plants affected by sudden-vegetation dieback in the Atlantic northeast (6). In our study, *Fusarium* spp. were found in only six leaf samples, all classified as damaged leaves. However, in all cases, their overall relative abundance was low (median relative abundance: 0.38% of fungal ITS reads). In five of the six cases, the identified fungus was *F. palustre*. Thus, although present and preferentially associated with damaged leaves, *Fusarium* spp. were not prevalent or highly abundant in our studied system. Notably, none of our studied marshes were experiencing widespread sudden-vegetation dieback.

### Environmental acquisition of the S. alterniflora phyllosphere microbiome

Salt marshes occupy a unique position at the transition zone between terrestrial and marine ecosystems. Consequently, the mechanisms by which marsh plants acquire their phyllosphere microbiome are also distinct. In better-studied plant systems, establishment of the phyllosphere microbiome can occur through multiple horizontal dispersal mechanisms, including rain, wind-mediated aerosol deposition, insects, and soil, as well as through seed-mediated vertical transmission (42). The relative contribution of each of these processes is largely unknown as it interacts with plant species and the degree of environmental exposure of leaves (43, 44). Because greenhouse plants in our study were propagated from non-sterilized botanical seeds, any vertically transmitted microbiota should have been retained during germination. The near absence of phyllosphere microbiota in greenhouse plants therefore indicates that the *S. alterniflora* phyllosphere microbiome is acquired predominantly through horizontal transmission from the environment. Beyond common dispersal mechanisms, tidal influence and daily inundation by saline tidal water in salt marshes may serve as an additional pathway for microbial acquisition. Further, our observation that halophilic microbes were predominant among the phyllosphere microbiota of *S. alterniflora* suggests that this ecological niche is defined in part by salt stress. Consequently, the most abundant prokaryotic and fungal genera in the *S. alterniflora* phyllosphere include taxa that thrive under extremely saline conditions and have previously been found in hypersaline environments such as salterns or salt flats, including bacteria from the *Larsenimonas*, *Pseudomaribius*, *Salinimonas*, and *Palleronia* genera (27, 45–48). Members of *Alternaria* have also been described as halotolerant fungi, having been isolated from sediments in sea salt fields (49). The finding of these taxa in saline marine sediments opens the possibility that sediment serves as a primary source for the phyllosphere microbiome, potentially mediated by microbial resuspension in tidal water. Insects may also contribute to phyllosphere microbial acquisition. *Candidatus* Uzinura, the most abundant prokaryote in natural marsh samples from the tall phenoype, is an obligate symbiont of scale insects (50), which are found on *S. alterniflora* leaves along the eastern U.S. coast (51). The detection of *Candidatus* Uzinura predominantly in the tall phenotype, which harbors the highest insect densities (52), suggests that insect activity may serve as a source of environmental microbes to the phyllosphere. Controlled experiments are needed to determine the relative contribution of sediment resuspension, tidal inundation, and insect activity to the assembly of the *S. alterniflora* phyllosphere microbiome.

### Implications for salt marsh restoration

The phyllosphere microbiome has been shown to outcompete pathogens, provide nutrients by nitrogen fixation, and detoxify methanol through methylotrophy (42). For instance, prior studies have shown that microbial depletion in the phyllosphere results in increased vulnerability to pathogenic infections (53, 54). Thus, based on our results, we suggest that inoculation of microbially depleted greenhouse leaves may help plants withstand pathogenic infection in both greenhouse and field settings. Early promising studies have shown that rhizosphere microbial inoculation in *S. alterniflora* can lead to increased biomass accumulation under high sediment salinity (55). Similarly, a microbial isolate retrieved from the phyllosphere of the mangrove plant *A. marina* has been shown to improve salinity stress tolerance in rice (31). However, to the best of our knowledge, inoculation of phyllosphere communities with beneficial microbes has not yet been tested in *S. alterniflora*. Further studies are needed to test and select microbial species, synthetic communities, or whole-microbiome inoculation strategies for the *S. alterniflora* phyllosphere. This plant is widely used in coastal restoration efforts, with individuals commonly grown in greenhouse settings and transplanted to the field as seedlings. Because the timing of species arrival strongly influences phyllosphere microbiome assembly (i.e., priority effects; (56)), our finding of microbial depletion in greenhouse leaves opens the possibility of successful inoculation with beneficial microbes prior to transplanting into marsh habitat, taking advantage of the largely unoccupied ecological niche on greenhouse leaves. By providing an *in situ* baseline of the *S. alterniflora* phyllosphere microbiome under contrasting health and environmental stress conditions, this study offers a benchmark for evaluating how closely greenhouse-based microbial inoculation outcomes replicate results observed in the field.

## Materials and Methods

### Natural marsh sampling

To study the phyllosphere microbiome in the natural habitat of *S. alterniflora*, we conducted leaf sampling within the state of Georgia during the July peak growth of *S. alterniflora* in 2021 and 2022. Leaf samples were collected from established transects on Sapelo Island and Skidaway Island, GA, where the sediment and root microbiomes of *S. alterniflora* have been previously studied (Figure 1A; for further details, see Rolando et al. (19)). Transects were established along a natural environmental stress gradient characterized by variation in sulfide concentration, salinity, and sediment redox conditions (Figure 1C). This gradient produces distinct phenotypes in *S. alterniflora*, with stressed sites supporting plants of the short phenotype (<100 cm) and less stressful sites supporting plants of the tall phenotype (>100 cm) (17). In total, 110 healthy and 80 damaged plants were collected from natural marshes in Georgia (Table 1). Only leaves from the top third of the plant were collected to avoid senescing leaves. Leaf health status was classified as healthy or damaged based on field observations alone. Damaged leaves presented one or more of the following: lesions, necrosis, spotting, or visible surface microbial growth, whereas healthy leaves lacked visible signs of tissue damage. Samples were collected using aseptic techniques and stored frozen until nucleic acid extraction.

### Collection of greenhouse plants

Leaf samples were collected from plants grown in two greenhouse facilities located in Savannah, GA, and Charleston, SC. Plants were propagated from botanical seeds collected from local marsh populations and stratified submerged in tap water at 4°C for a minimum of 8 weeks prior to germination in freshwater. Following germination, seedlings were transplanted into either a 2:1 topsoil:sand mixture (Savannah) or commercial potting mix (Charleston). Plants were 3–5 months old at the time of sampling. Leaf samples were collected aseptically and stored at −80°C until DNA extraction.

### Nucleic acids Extraction and Amplicon Sequencing

Leaf tissue was ground in liquid nitrogen, followed by DNA extraction using the DNeasy PlantPro Kit (Qiagen, Valencia, CA) according to the manufacturer’s protocol. DNA was quantified using the Qubit HS assay (Invitrogen, Carlsbad, CA), and purity was measured using the NanoDrop 2000/2000c spectrophotometer (Thermo Fisher Scientific, Waltham, MA, USA). The V4 region of the 16S rRNA gene was amplified to characterize the prokaryotic community using primers 515F (5′–GTGYCAGCMGCCGCGGTAA) and 806R (5′– GGACTACNVGGGTWTCTAAT) (57, 58). Fungal communities were characterized by amplifying the ITS region using primers 5.8S-Fun (5′–AACTTTYRRCAAYGGATCWCT) and ITS4-Fun (5′– AGCCTCCGCTTATTGATATGCTTAART) (59). Peptide nucleic acid (PNA) clamps were used to reduce plastid and mitochondrial amplification during 16S rRNA gene amplification, as described in Lundberg et al. (60). PCR products were barcoded and sequenced on an Illumina MiSeq platform (v3 kit, PE300) at the Molecular Evolution Core at the Georgia Institute of Technology.

### Bioinformatic and Statistical Analysis

Amplicon sequences were analyzed for both prokaryotic and fungal communities. Cutadapt v.2.0 was used to remove primers from raw FASTQ files (61). DADA2 v.1.29.0 was used to infer amplicon sequence variants (ASVs) from quality-filtered reads (62). Chimeras were removed using the removeBimeraDenovo function in the DADA2 package. Taxonomy was assigned to 16S rRNA gene amplicon sequences using the SILVA SSU rRNA database (version 138.2; (63)) and to ITS amplicon sequences using the UNITE database (version 9.0; (64)). All ASVs from unclassified kingdoms were excluded from further analysis.

Prokaryotic relative abundance was calculated as the percentage of sequence reads originating from prokaryotic ASVs out of the total number of 16S rRNA gene sequence reads, including plant organelle 16S rRNA gene ASVs. Similarly, fungal relative abundance was calculated as the percentage of sequence reads originating from fungal ITS ASVs out of the total number of ITS sequence reads, including plant ITS ASVs. The Shannon diversity index was used to characterize the local diversity of the *S. alterniflora* phyllosphere microbiome and was calculated using the phyloseq v.1.44.0 package (65). The Shannon diversity index was calculated on microbial amplicon datasets after removing plant-associated reads. A two-way ANOVA was performed to investigate the interactive effects of plant health status (i.e., healthy and damaged) and *Spartina* phenotype (i.e., short and tall) on Shannon diversity for both 16S rRNA gene and ITS datasets, followed by Tukey’s honestly significant difference (HSD) post hoc tests when significance was detected. Because relative abundance data did not meet normality assumptions, they were analyzed using non-parametric Kruskal–Wallis tests followed by Dunn’s multiple comparison tests. For greenhouse plants, the same models were used to test for differences in Shannon diversity and relative abundance relative to healthy plants from both tall and short natural phenotypes.

Beta diversity was assessed on microbial reads only by calculating the Bray-Curtis dissimilarity matrix using the ordinate function in the phyloseq package. Nonmetric multidimensional scaling (NMDS) ordinations were then performed for visualization. For natural marsh plants, the variance of the Bray-Curtis dissimilarity matrix was partitioned based on *S. alterniflora* phenotype, health status, and site location (Sapelo – GCE06, Sapelo – Sentinel, and Skidaway Island) using permutational multivariate analysis of variance (PERMANOVA) with 999 permutations. When comparing greenhouse plants to healthy natural plants, PERMANOVA was performed to partition the Bray-Curtis dissimilarity matrix by *Spartina* phenotype (greenhouse, tall, and short) and site location (Sapelo – GCE06, Sapelo – Sentinel, Skidaway Island, GSU Greenhouse, and South Carolina Greenhouse). PERMANOVA was conducted using the adonis2 function in the vegan v.2.6-4 package.

Finally, the Microbiome Multivariable Associations with Linear Models 2 (MaAsLin2; (24)) R package was used to identify microorganisms that were differentially abundant under contrasting plant health statuses, *S. alterniflora* phenotypes, and growth conditions (greenhouse or natural salt marsh). Amplicon data were first normalized to relative abundance using the full set of reads, including plant-derived organelle 16S rRNA gene and plant ITS sequences, so that microbial abundances were expressed relative to total plant-associated DNA. After relative abundances were calculated, plant-derived reads were removed from the dataset prior to MaAsLin2 analysis, as the objective was to identify differentially abundant microbial taxa rather than variation in plant sequences. Multiple-testing correction was performed using the Benjamini–Hochberg (BH) method. Three separate models were run in MaAsLin2 to address distinct questions. First, to identify microorganisms that were more abundant in damaged leaves, only natural marsh samples were analyzed, with plant health status and *Spartina* phenotype as fixed factors. Second, a subset of only healthy plants was analyzed with *Spartina* phenotype (i.e., greenhouse, short, and tall) as a fixed factor to identify how greenhouse plants differed in their phyllosphere microbiome relative to healthy short and tall plants from natural marshes. Finally, only damaged leaves from natural marsh plants were analyzed with *Spartina* phenotype (i.e., short and tall) as the fixed factor to identify differences in microbial ASV abundance in damaged plants experiencing contrasting levels of environmental stress.

## ACKNOWLEDGMENTS

This publication is supported in part by the U.S. National Science Foundation via the Georgia Coastal Ecosystems LTER program (OCE-1832178, and OCE-2425396) and an institutional grant NA24OARX417C0160-T1-01 to the Georgia Sea Grant College Program from the National Sea Grant Office under the National Oceanic and Atmospheric Administration, U.S. Department of Commerce. All views, opinions, findings, conclusions, and recommendations expressed in this material are those of the author(s) and do not necessarily reflect the opinions of the Georgia Sea Grant College Program or the National Oceanic and Atmospheric Administration. This is contribution № 1,141 of the University of Georgia Marine Institute. J.L.R.: writing—original draft, data curation, visualization, investigation, methodology, formal analysis, conceptualization. A.L.C.: writing—original draft, methodology, investigation. M.H.: resources, investigation. H.J.: resources, investigation. J.E.K.: writing—original draft, methodology, investigation, conceptualization, supervision, project administration, funding acquisition.

## DATA AVAILABILITY

Raw SSU rRNA gene and ITS amplicon sequences have been deposited in the BioProject database (http://ncbi.nlm.nih.gov/bioproject) under accession PRJNA1423775.

## REFERENCES

1. Barbier EB, Hacker SD, Kennedy C, Koch E, Stier AC, Silliman BR. 2011. The value of estuarine and coastal ecosystem services. Ecological Monographs 81:169–193.

2. Alber M, Swenson EM, Adamowicz SC, Mendelssohn IA. 2008. Salt Marsh Dieback: An overview of recent events in the US. Estuarine, Coastal and Shelf Science 80:1–11.

3. Gu J, Luo M, Zhang X, Christakos G, Agusti S, Duarte CM, Wu J. 2018. Losses of salt marsh in China: Trends, threats and management. Estuarine, Coastal and Shelf Science 214:98–109.

4. Borges FO, Santos CP, Paula JR, Mateos-Naranjo E, Redondo-Gomez S, Adams JB, Caçador I, Fonseca VF, Reis-Santos P, Duarte B, Rosa R. 2021. Invasion and Extirpation Potential of Native and Invasive Spartina Species Under Climate Change. Front Mar Sci 8:696333.

5. Elmer WH. 2014. A Tripartite Interaction Between *Spartina alterniflora*, *Fusarium palustre*, and the Purple Marsh Crab (*Sesarma reticulatum*) Contributes to Sudden Vegetation Dieback of Salt Marshes in New England. Phytopathology® 104:1070–1077.

6. Elmer WH, Marra RE. 2011. New species of *Fusarium* associated with dieback of *Spartina alterniflora* in Atlantic salt marshes. Mycologia 103:806–819.

7. Elmer WH, Useman S, Schneider RW, Marra RE, LaMondia JA, Mendelssohn IA, Jiménez-Gasco MM, Caruso FL. 2013. Sudden vegetation dieback in Atlantic and Gulf coast salt marshes. Plant Disease 97:436–445.

8. Daleo P, Alberti J, Pascual J, Iribarne O. 2013. Nutrients and Abiotic Stress Interact to Control Ergot Plant Disease in a SW Atlantic Salt Marsh. Estuaries and Coasts 36:1093– 1097.

9. Kaur R, Knott C, Aime MC. 2010. First Report of Rust Disease Caused by Puccinia sparganioides on Spartina alterniflora in Louisiana. Plant Disease 94:636–636.

10. Pfeilmeier S, Werz A, Ote M, Bortfeld-Miller M, Kirner P, Keppler A, Hemmerle L, Gäbelein CG, Petti GC, Wolf S, Pestalozzi CM, Vorholt JA. 2024. Leaf microbiome dysbiosis triggered by T2SS-dependent enzyme secretion from opportunistic Xanthomonas pathogens. Nat Microbiol 9:136–149.

11. Runge P, Ventura F, Kemen E, Stam R. 2023. Distinct Phyllosphere Microbiome of Wild Tomato Species in Central Peru upon Dysbiosis. Microb Ecol 85:168–183.

12. Krishnappa C, Sahu KP, Ashajyothi M, Kumar M, Reddy B, Kumar A. 2025. Dysbiosis of the rice leaf phyllomicrobiome induced by Magnaporthe oryzae infection: evidence from metabarcoding and microbiome imprinting. Int Microbiol 28:2377–2389.

13. Chen T, Nomura K, Wang X, Sohrabi R, Xu J, Yao L, Paasch BC, Ma L, Kremer J, Cheng Y, Zhang L, Wang N, Wang E, Xin X-F, He SY. 2020. A plant genetic network for preventing dysbiosis in the phyllosphere. Nature 580:653–657.

14. Tang L, Gao Y, Li B, Wang Q, Wang C-H, Zhao B. 2014. *Spartina alterniflora* with high tolerance to salt stress changes vegetation pattern by outcompeting native species. Ecosphere 5:1–18.

15. Bradley PM, Morris JT. 1991. Relative importance of ion exclusion, secretion and accumulation in Spartina alterniflora loisel. Journal of Experimental Botany 42:1525–1532.

16. Mendelssohn IA, Morris JT. 2002. Eco-Physiological Controls on the Productivity of Spartina Alterniflora Loisel, p. 59–80. In Concepts and Controversies in Tidal Marsh Ecology. Springer Netherlands, Dordrecht.

17. Dai T, Wiegert RG. 1996. Ramet population dynamics and net aerial primary productivity of Spartina Alterniflora. Ecology 77:276–288.

18. Thomas F, Giblin AE, Cardon ZG, Sievert SM. 2014. Rhizosphere heterogeneity shapes abundance and activity of sulfur-oxidizing bacteria in vegetated salt marsh sediments. Frontiers in Microbiology 5:1–14.

19. Rolando JL, Kolton M, Song T, Kostka JE. 2022. The core root microbiome of Spartina alterniflora is predominated by sulfur-oxidizing and sulfate-reducing bacteria in Georgia salt marshes, USA. Microbiome 10:1–17.

20. Du Plessis I, Snyder H, Calder R, Rolando JL, Kostka JE, Weitz JS, Dominguez-Mirazo M. 2025. Viral community diversity in the rhizosphere of the foundation salt marsh plant *Spartina alterniflora*. mSphere 10:e00234–25.

21. Buchan A, Newell SY, Butler M, Biers EJ, Hollibaugh JT, Moran MA. 2003. Dynamics of Bacterial and Fungal Communities on Decaying Salt Marsh Grass. Appl Environ Microbiol 69:6676–6687.

22. Song Z, Huang Y, Liu Q, Hu X. 2022. Discovering the Characteristics of Community Structures and Functional Properties of Epiphytic Bacteria on Spartina alterniflora in the Coastal Salt Marsh Area. JMSE 10:1981.

23. Majeed A, Liu J, Knight AJ, Pajerowska-Mukhtar KM, Mukhtar MS. 2024. Bacterial Communities Associated with the Leaves and the Roots of Salt Marsh Plants of Bayfront Beach, Mobile, Alabama, USA. Microorganisms 12:1595.

24. Mallick H, Rahnavard A, McIver LJ, Ma S, Zhang Y, Nguyen LH, Tickle TL, Weingart G, Ren B, Schwager EH, Chatterjee S, Thompson KN, Wilkinson JE, Subramanian A, Lu Y, Waldron L, Paulson JN, Franzosa EA, Bravo HC, Huttenhower C. 2021. Multivariable association discovery in population-scale meta-omics studies. PLoS Comput Biol 17:e1009442.

25. Antunes A, Eder W, Fareleira P, Santos H, Huber R. 2003. Salinisphaera shabanensis gen. nov., sp. nov., a novel, moderately halophilic bacterium from the brine–seawater interface of the Shaban Deep, Red Sea. Extremophiles 7:29–34.

26. Koblížek M, Béjà O, Bidigare RR, Christensen S, Benitez-Nelson B, Vetriani C, Kolber MK, Falkowski PG, Kolber ZS. 2003. Isolation and characterization of Erythrobacter sp. strains from the upper ocean. Arch Microbiol 180:327–338.

27. Yoon J-H, Kang S-J, Lee S-Y. 2012. Salinimonas lutimaris sp. nov., a polysaccharide- degrading bacterium isolated from a tidal flat. Antonie van Leeuwenhoek 101:803–810.

28. Navarro-Torre S, Carro L, Rodríguez-Llorente ID, Pajuelo E, Caviedes MÁ, Igual JM, Redondo-Gómez S, Camacho M, Klenk H-P, Montero-Calasanz MDC. 2018. Kushneria phyllosphaerae sp. nov. and Kushneria endophytica sp. nov., plant growth promoting endophytes isolated from the halophyte plant Arthrocnemum macrostachyum. International Journal of Systematic and Evolutionary Microbiology 68:2800–2806.

29. Fidalgo C, Proença DN, Morais PV, Henriques I, Alves A. 2019. The endosphere of the salt marsh plant Halimione portulacoides is a diversity hotspot for the genus Salinicola: description of five novel species Salinicola halimionae sp. nov., Salinicola aestuarinus sp. nov., Salinicola endophyticus sp. nov., Salinicola halophyticus sp. nov. and Salinicola lusitanus sp. nov. International Journal of Systematic and Evolutionary Microbiology 69:46–62.

30. Vorholt JA. 2012. Microbial life in the phyllosphere. Nat Rev Microbiol 10:828–840.

31. Yang X, Dai Z, Yuan R, Guo Z, Xi H, He Z, Wei M. 2023. Effects of Salinity on Assembly Characteristics and Function of Microbial Communities in the Phyllosphere and Rhizosphere of Salt-Tolerant *Avicennia marina* Mangrove Species. Microbiol Spectr 11:e03000–22.

32. Bergbauer M, Newell SY. 1992. Contribution to lignocellulose degradation and DOC formation from a salt marsh macrophyte by the aseomycete Phaeosphaeria spartinicola. FEMS Microbiology Letters 86:341–347.

33. Petersen C, Round JL. 2014. Defining dysbiosis and its influence on host immunity and disease. Cell Microbiol 16:1024–1033.

34. Hutchinson GE. 1959. Homage to Santa Rosalia or Why Are There So Many Kinds of Animals? The American Naturalist 93:145–159.

35. Yu XA, McLean C, Hehemann J-H, Angeles-Albores D, Wu F, Muszyński A, Corzett CH, Azadi P, Kujawinski EB, Alm EJ, Polz MF. 2024. Low-level resource partitioning supports coexistence among functionally redundant bacteria during successional dynamics. The ISME Journal 18:wrad013.

36. Daleo P, Silliman B, Alberti J, Escapa M, Canepuccia A, Peña N, Iribarne O. 2009. Grazer facilitation of fungal infection and the control of plant growth in south-western Atlantic salt marshes. Journal of Ecology 97:781–787.

37. Bärlocher F, Newell SY. 1994. Growth of the salt marsh periwinkleLittoraria irrorata on fungal and cordgrass diets. Mar Biol 118:109–114.

38. Walker AK, Campbell J. 2010. Marine fungal diversity: a comparison of natural and created salt marshes of the north-central Gulf of Mexico. Mycologia 102:513–521.

39. Lyons JI, Alber M, Hollibaugh JT. 2010. Ascomycete fungal communities associated with early decaying leaves of Spartina spp. from central California estuaries. Oecologia 162:435–442.

40. Elmer WH. 2016. Pathogenic Microfungi Associated with Spartina in Salt Marshes 615– 630.

41. Sieg RD, Wolfe K, Willey D, Ortiz-Santiago V, Kubanek J. 2013. Chemical defenses against herbivores and fungi limit establishment of fungal farms on salt marsh angiosperms. Journal of Experimental Marine Biology and Ecology 446:122–130.

42. Koskella B. 2020. The phyllosphere. Current Biology 30:R1143–R1146.

43. Abdelfattah A, Tack AJM, Lobato C, Wassermann B, Berg G. 2023. From seed to seed: the role of microbial inheritance in the assembly of the plant microbiome. Trends in Microbiology 31:346–355.

44. Chesneau G, Laroche B, Préveaux A, Marais C, Briand M, Marolleau B, Simonin M, Barret M. 2022. Single Seed Microbiota: Assembly and Transmission from Parent Plant to Seedling. mBio 13:e01648–22.

45. Jeon CO, Lim J-M, Park D-J, Kim C-J. 2005. Salinimonas chungwhensis gen. nov., sp. nov., a moderately halophilic bacterium from a solar saltern in Korea. International Journal of Systematic and Evolutionary Microbiology 55:239–243.

46. Kim Y, Kim J-H, Lee KC, Lee J-S, Kim W. 2015. Palleronia soli sp. nov., isolated from a soil sample on reclaimed tidal land, and emended description of the genus Palleronia. International Journal of Systematic and Evolutionary Microbiology 65:2516–2521.

47. Xia Z-J, Wu H-Z, Cui C-X, Chen Q, Zhao G-Y, Wang H-X, Dai M-X. 2016. Larsenimonas suaedae sp. nov., a moderately halophilic, endophytic bacterium isolated from the euhalophyte Suaeda salsa. International Journal of Systematic and Evolutionary Microbiology 66:2952–2958.

48. Zhang H, Wang H, Cao L, Chen H, Wang M, Lian C, Zhong Z, Li C. 2020. Salinimonas iocasae sp. nov., a halophilic bacterium isolated from a polychaete tube in a hydrothermal field. International Journal of Systematic and Evolutionary Microbiology 70:3899–3904.

49. Wang W, Wang Y, Tao H, Peng X, Liu P, Zhu W. 2009. Cerebrosides of the Halotolerant Fungus *Alternaria raphani* Isolated from a Sea Salt Field. J Nat Prod 72:1695–1698.

50. Gruwell ME, Flarhety M, Dittmar K. 2012. Distribution of the Primary Endosymbiont (Candidatus Uzinura Diaspidicola) Within Host Insects from the Scale Insect Family Diaspididae. Insects 3:262–269.

51. Japoshvili G, Russell E. 2012. A New Parasitization Record of *Haliaspis spartinae* (Diaspididae) and *Encarsia ellisvillensis* sp. nov. (Chalcidoidea: Aphelinidae) from the United States. Annals of the Entomological Society of America 105:493–497.

52. Gaeta JW, Kornis MS. 2011. Stem Borer Frequency and Composition in Healthy Spartina alterniflora (smooth cordgrass) and Dieback Zones in a Southern Atlantic Coast Salt Marsh. Estuaries and Coasts 34:1078–1083.

53. Berg M, Koskella B. 2018. Nutrient- and Dose-Dependent Microbiome-Mediated Protection against a Plant Pathogen. Current Biology 28:2487–2492.e3.

54. Paasch BC, Sohrabi R, Kremer JM, Nomura K, Cheng YT, Martz J, Kvitko B, Tiedje JM, He SY. 2023. A critical role of a eubiotic microbiota in gating proper immunocompetence in Arabidopsis. Nat Plants 9:1468–1480.

55. Bledsoe R, Boopathy R. 2016. Bioaugmentation of microbes to restore coastal wetland plants to protect land from coastal erosion. International Biodeterioration and Biodegradation 113:155–160.

56. Trivedi P, Leach JE, Tringe SG, Sa T, Singh BK. 2020. Plant–microbiome interactions: from community assembly to plant health. Nature Reviews Microbiology 18:607–621.

57. Apprill A, McNally S, Parsons R, Weber L. 2015. Minor revision to V4 region SSU rRNA 806R gene primer greatly increases detection of SAR11 bacterioplankton. Aquat Microb Ecol 75:129–137.

58. Parada AE, Needham DM, Fuhrman JA. 2016. Every base matters: assessing small subunit RRNA primers for marine microbiomes with mock communities, time series and global field samples. Environmental Microbiology 18:1403–1414.

59. Taylor DL, Walters WA, Lennon NJ, Bochicchio J, Krohn A, Caporaso JG, Pennanen T. 2016. Accurate Estimation of Fungal Diversity and Abundance through Improved Lineage- Specific Primers Optimized for Illumina Amplicon Sequencing. Appl Environ Microbiol 82:7217–7226.

60. Lundberg DS, Yourstone S, Mieczkowski P, Jones CD, Dangl JL. 2013. Practical innovations for high-throughput amplicon sequencing. Nat Methods 10:999–1002.

61. Martin M. 2011. Cutadapt removes adapter sequences from high-throughput sequencing reads. EMBnet journal 17:10–12.

62. Callahan BJ, McMurdie PJ, Rosen MJ, Han AW, Johnson AJA, Holmes SP. 2016. DADA2: High-resolution sample inference from Illumina amplicon data. Nat Methods 13:581–583.

63. Chuvochina M, Gerken J, Frentrup M, Sandikci Y, Goldmann R, Freese HM, Göker M, Sikorski J, Yarza P, Quast C, Peplies J, Glöckner FO, Reimer LC. 2026. SILVA in 2026: a global core biodata resource for rRNA within the DSMZ digital diversity. Nucleic Acids Research 54:D334–D341.

64. Abarenkov K, Zirk A, Piirmann T, Pöhönen R, Ivanov F, Nilsson RH, Kõljalg. 2023. Full UNITE+INSD dataset for eukaryotes (18.07.2023). UNITE Community.

65. McMurdie PJ, Holmes S. 2013. phyloseq: An R Package for Reproducible Interactive Analysis and Graphics of Microbiome Census Data. PLoS ONE 8:e61217.

