## Supplementary Figures and Tables for "Leaf Damage Is Associated with Alteration of Microbiome Composition and Abundance in the *Spartina alterniflora* Phyllosphere"

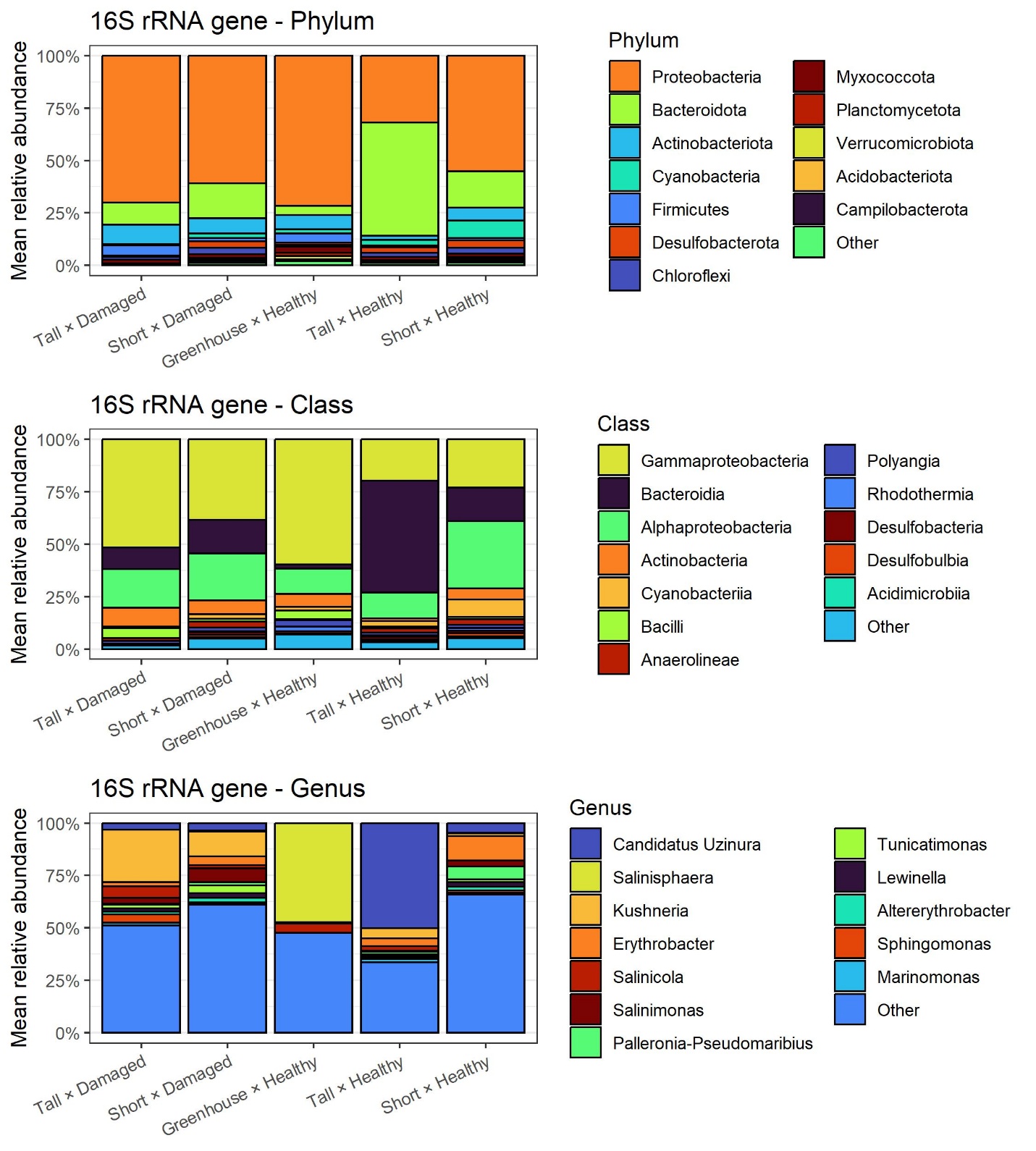

**Figure S1. Prokaryotic composition of the *Spartina alterniflora* phyllosphere microbiome**

Stacked bar plots representing the mean relative abundance of prokaryotic taxa detected by 16S rRNA gene amplicon sequencing in the phyllosphere of *Spartina alterniflora*. Plant organelle 16S rRNA gene reads were discarded before plotting. Taxonomic composition is shown at the (A) phylum, (B) class, and (C) genus levels. Samples are grouped by plant phenotype (Tall vs. Short), plant health status (Healthy vs. Damaged), and greenhouse-grown plants where indicated. Only the top 12 most abundant taxa at each rank are shown individually, while the remaining low-abundance taxa are grouped into the “Other” category.

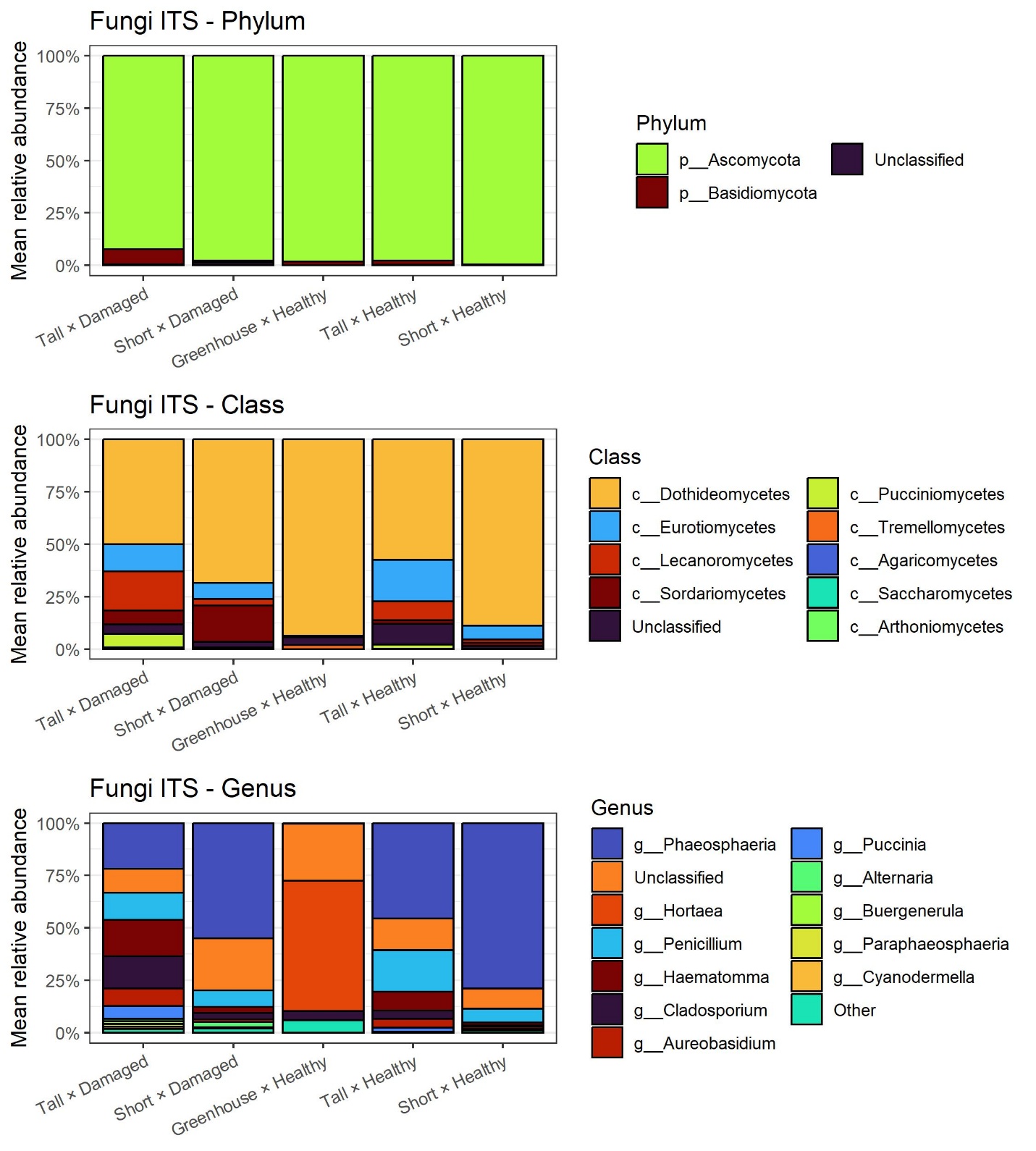

**Figure S2. Fungal composition of the *Spartina alterniflora* phyllosphere microbiome**

Stacked bar plots representing the mean relative abundance of fungal taxa detected by ITS region amplicon sequencing in the phyllosphere of *Spartina alterniflora*. Plant ITS reads were discarded before plotting. Taxonomic composition is shown at the (A) phylum, (B) class, and (C) genus levels. Samples are grouped by plant phenotype (Tall vs. Short), plant health status (Healthy vs. Damaged), and greenhouse-grown plants where indicated. Only the top 12 most abundant taxa at each rank are shown individually, while the remaining low-abundance taxa are grouped into the “Other” category.

| **Table S1:** Two-way analysis of variance (ANOVA) of the Shannon diversity index as explained by Spartina phenotype, plant health status, and its interaction | | | | |
| --- | --- | --- | --- | --- |
|  | 16S rRNA | | ITS | |
| Factor | F-value | p-value | F-value | p-value |
| Spartina Phenotype | 76.67 | 2.7E-15 | 2.17 | 0.143 |
| Plant Health | 5.70 | 0.018 | 7.33 | 0.007 |
| Interaction | 9.33 | 0.003 | 0.04 | 0.850 |
